# SARS-CoV-2 shedding dynamics in human respiratory tract

**DOI:** 10.1101/2024.07.09.602697

**Authors:** Xiangxing Jin, Lili Ren, Xianwen Ren, Jianwei Wang

## Abstract

It is crucial to understand how Severe Acute Respiratory Syndrome Coronavirus 2 (SARS-CoV-2) sheds in human respiratory tract, but this question is still elusive to understand due to technical limitations. Here we integrated published human metagenomic data of SARS-CoV-2 and developed a novel algorithm named as RedeCoronaVS to systematically dissect SARS-CoV-2 shedding modes with single-cell data as reference. We identify that SARS-CoV-2 particles are the dominant mode of viral shedding in the very early infection phase (≤24 hours after hospitalization). Within the first week after hospitalization, SARS-CoV-2 replicons within host cells dominate SARS-CoV-2 shedding together with viral particles. One week later, viral fragments become the dominant mode in patients with mild or moderate symptoms, but viral replicons still dominate in some patients with severe symptoms. In addition to epithelial cells, SARS-CoV-2 replicons in neutrophils, macrophages, and plasma cells also show important roles and are associate with sampling time and disease severity.

## Introduction

Coronavirus Disease 2019 (COVID-19), representing a newly emerging infectious disease, is a respiratory illness known for rapid transmission, acute onset, diverse modes of transmission, and the potential for multi-organ infections.^1^ Despite relaxed control policies in many countries, the ongoing emergence of new SARS-CoV-2 variants poses a risk of recurrent infections for susceptible populations.^2^ Therefore, understanding the viral shedding patterns of COVID-19 patients is important for formulating precise control strategies. Currently, research on methods for detecting viral shedding in COVID-19 patients mainly falls into three categories: quantitative reverse transcription polymerase chain reaction (qRT-PCR) and droplet digital polymerase chain reaction (ddPCR) for measuring viral RNA copies; antigen-detecting (rapid) diagnostic tests targeting viral proteins, primarily the spike protein; and methods for measuring infectious viral particles, such as virus isolation and plaque assays or 50% tissue culture infectious dose (TCID_50_)^3–5^ However, conventional methods measuring viral RNA are limited to quantification, rather than dissecting the shedding patterns.^3^ The viral load typically represents the number of viral RNA copies, but it only quantifies the viral genome fragments and does not represent infectious viral particles.^6^ Methods customized for viral particle analysis, such as electron microscopy or plaque-forming assays, can offer compelling evidence but are hard to be applied to large number of samples due to high cost and biosafety requirements.^7,8^ Therefore, there is an urgent need to develop new techniques to understand the patterns of SARS-CoV-2 shedding based on large cohorts.

Metatranscriptomic sequencing is a direct and unbiased method for acquiring comprehensive transcriptome sequencing data from samples, which has been widely employed in current microbiological research and is highly efficient for virus discovery.^9,10^ Recent studies have used this approach to reveal the microbial microenvironment in the respiratory tract of COVID-19 patients and exploring the relationship between the microbiome and transcriptome.^11^ We have observed that metatranscriptomic sequencing techniques, covering the entire transcriptome of SARS-CoV-2, offer a burgeoning research perspective on viral shedding in COVID-19 patients. However, there is still a lack of tools to integrate these two aspects.

Here, we gathered a total of 301 samples with metatranscriptomic sequencing data from the respiratory tracts of COVID-19 patients, encompassing diverse nationalities, ethnicities, infection stages, disease severities, and sample types. These samples, which includes comprehensive clinical information of the patients, stands as the most comprehensive to date. Based on single-cell sequencing and metatranscriptomic sequencing data, we developed a digital analytical method named as RedeCoronaVS to dissect the patterns of SARS-CoV-2 shedding in the respiratory tract of COVID-19 patients. We categorized viral shedding in the respiratory tract into three types of patterns: viral replicons (intracellular), viral particles (extracellular), and viral fragments. Unsupervised machine learning results based on non-negative matrix factorization prove our categorization. Then we developed RedeCoronaVS, which accurately determines the abundance of various shedding patterns in individual patients through the viral RNA data, enhancing our understanding of viral shedding in COVID-19 patients. RedeCoronaVS revealed correlations between different patterns of viral shedding and clinical features such as sampling site, days after hospitalization, and the severity of COVID-19. The pattern of viral shedding in COVID-19 patients varies depending on the sampling time during hospitalization. The types of viral replicons are associated with the clinical status of patients. Neutrophil and macrophage viral replicons tend to shed in the early stages of infection (≤24 hours after hospitalization), while plasma cell viral replicons predominate in the respiratory tract of severe COVID-19 patients. Our integrative analysis, the newly developed algorithms, and the findings are expected to provide important insights into the viral shedding dynamics of COVID-19.

## Results

### Comparisons of the subgenomic transcription of SARS-CoV-2 among different cell types

As a typical coronavirus, SARS-CoV-2 exhibits ‘discontinuous transcription’ (also known as subgenomic transcription) within host cells, greatly enhancing transcription and translation efficiency (Figure 1A).^12,13^ While many studies have established the multi-tissue and multi-cell type tropism of SARS-CoV-2, it is unclear whether the virus exhibits different subgenomic transcription patterns in various cell types.^14,15^ To address this, we gathered data from previous study utilizing single-cell RNA sequencing to detect viral genes in different cell types infected by SARS-CoV-2.^16^ Given that ciliated cells are considered the primary host cells for SARS-CoV-2,^17,18^ we set the pattern of viral transcription in ciliated cells as a standard template and compared it with other cell types. Subsequent correlation analysis indicates slight heterogeneity in viral transcription among cell types, for example, differences in epithelial cells are mainly in the *E* and *ORF7b* genes (Figure 1B), while differences between ciliated cells and other cell types such as myeloid cells and lymphocytes are mainly in the *N* gene (Figure 1C-1D). This suggests the potential to distinguish infected cell types based on the detection rate of SARS-CoV-2.

**Figure 1.**
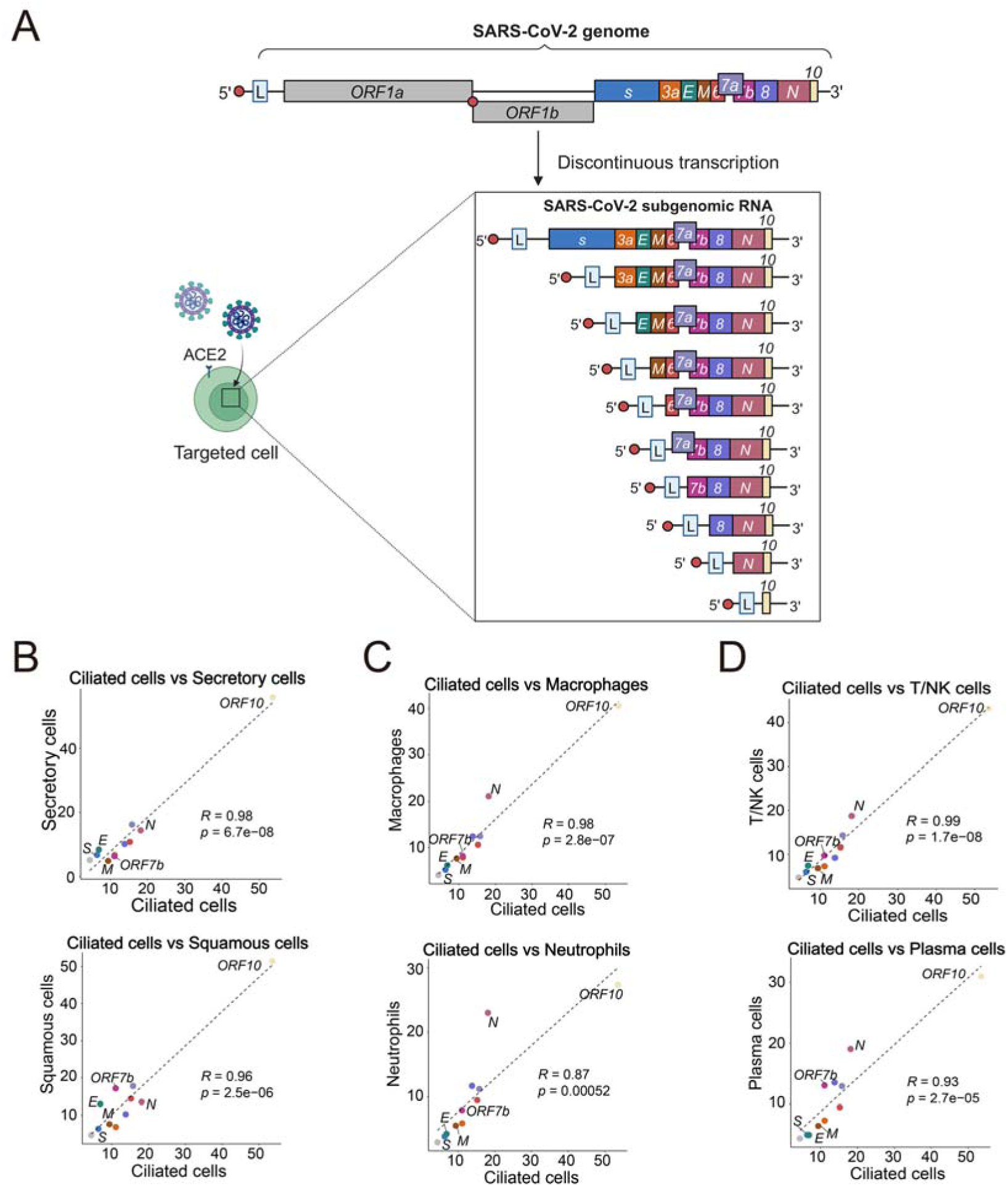
The subgenomic transcription patterns of SARS-CoV-2 in target cells. (A) Schematic representation of the subgenomic transcription landscape of SARS-CoV-2 within host cells. Created by BioRender.com. (B) Plot of the relationship between the SARS-CoV-2 detection rate in ciliated cells and other epithelial cells. (Pearson correlation, dotted line, the square root transformation was applied to the detection rate.). The viral detection rate is defined as the ratio of cells with positive viral genes in a specific cell type to the total number of cells of that type, with subsequent gene length normalization. (C) Plot of the relationship between the SARS-CoV-2 detection rate in ciliated cells and myeloid cells. (D) Plot of the relationship between the SARS-CoV-2 detection rate in ciliated cells and lymphocytes.

### Metatranscriptomic data of SARS-CoV-2 deviate subgenomic transcription

Next, we collected 301 respiratory samples with metatranscriptomic data available from three previously published cohorts (STAR Methods). To explore the gene expression pattern of SARS-CoV-2 in bulk RNA sequencing data and its association with single-cell RNA sequencing data, we conducted an analysis of the distribution of viral RNA-seq data (normalized by Transcripts Per Kilobase Million) across the mean values of various genes (Figure 2A-2D, Figure S1A). Furthermore, we applied a correlation analysis between the single cell level viral RNA detection rate in SARS-CoV-2 infected ciliated cells and viral RNA-seq data (Figure 2E-2H, Figure S1B). In particular, the metatranscriptomic data of early hospitalization COVID-19 patients were characterized by close proximity to the rate of viral detection at the single-cell level, indicative of active viral replication and transmission (Figure 2A, 2E).^19,20^ Results from metatranscriptomic data from other cohorts differed from those from early hospitalization in having high *ORF7b* abundance and low correlation with single-cell level viral RNA detection, suggesting that SARS-CoV-2 viral metatranscriptome sequencing data do not fully align with the discontinuous transcription pattern of the virus within cells (Figure 2B-D, Figure 2F-H). The lower respiratory source samples were consistent with the results of the upper respiratory samples (Figure S1A-S1B).

**Figure 2.**
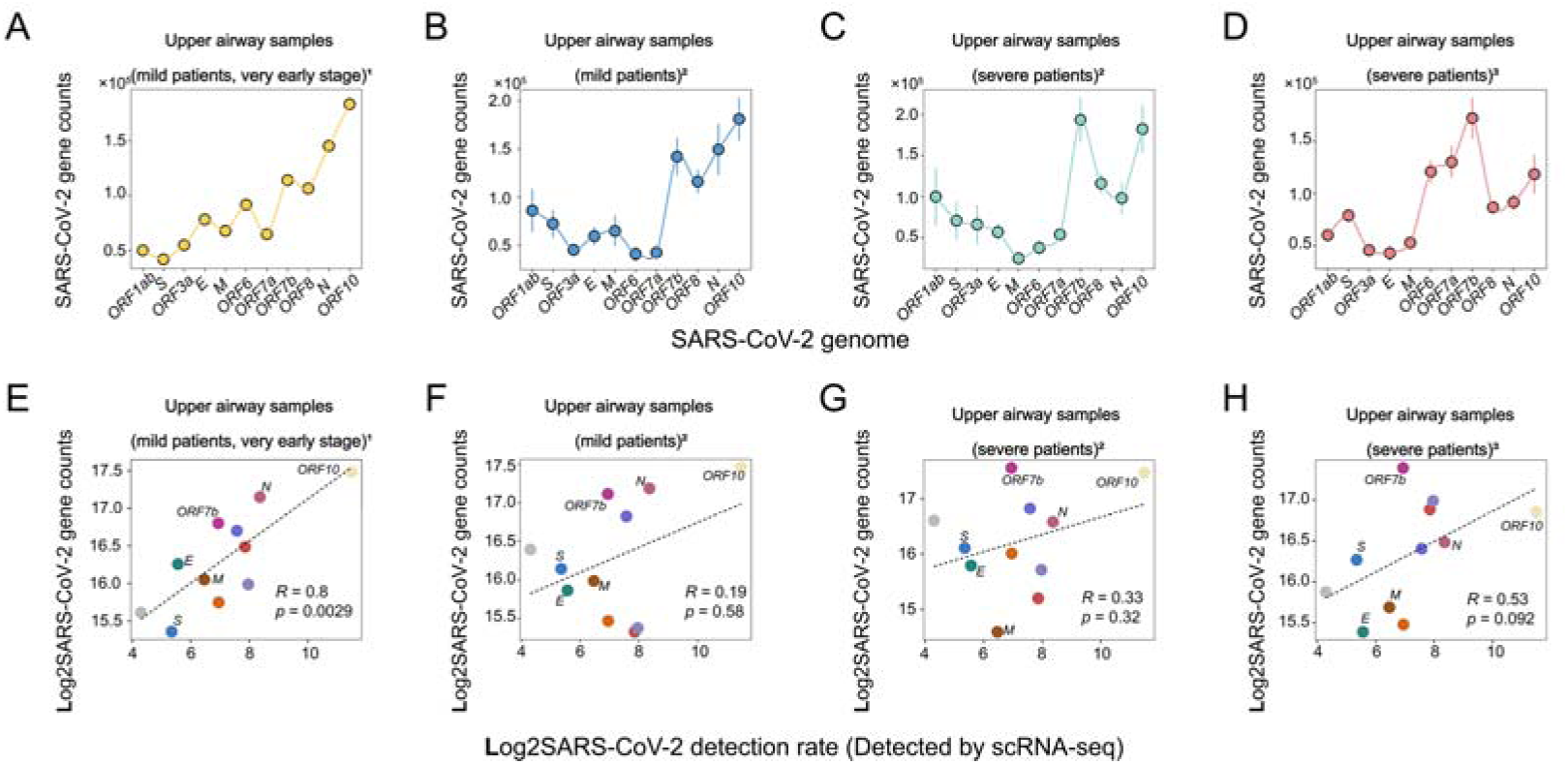
Associations between the transcription patterns of individual genes of SARS-CoV-2 in metatranscriptome data and their correlation with subgenomic transcription patterns. (A-D) Line graphs depicting the mean distribution of various viral genes in metatranscriptome samples. (Viral gene reads were normalized by TPM; standard error of the samples is indicated.). (E-H) Plots of the relationship between the normalized mean expression of SARS-CoV-2 viral genes in metatranscriptome samples across different types and the single-cell level detection rate of SARS-CoV-2 in ciliated cells. (Pearson correlation; dotted line, the logarithmic transformation was applied to viral detection rate.). The type and source of the samples are labeled. 1: Data from Rajagopala, Seesandra V et al. 2: Data from Ren, Lili et al. 3: Data from Sulaiman, Imran et al.

### Unsupervised dissection of SARS-CoV-2 shedding patterns in metatranscriptomic data

Based on the former results, we assuming that there are other potential confounding factors in the SARS-CoV-2 metatranscriptomic data, Non-Negative Matrix Factorization (NMF) was employed to identify the distribution characteristics of viral genes, facilitating the recognition of viral shedding patterns (STAR methods). When specifying three categories, the viral RNA distribution in the classification results is depicted in the figure (Figure 3A-3D, Figure S1C; Table S1).

**Figure 3.**
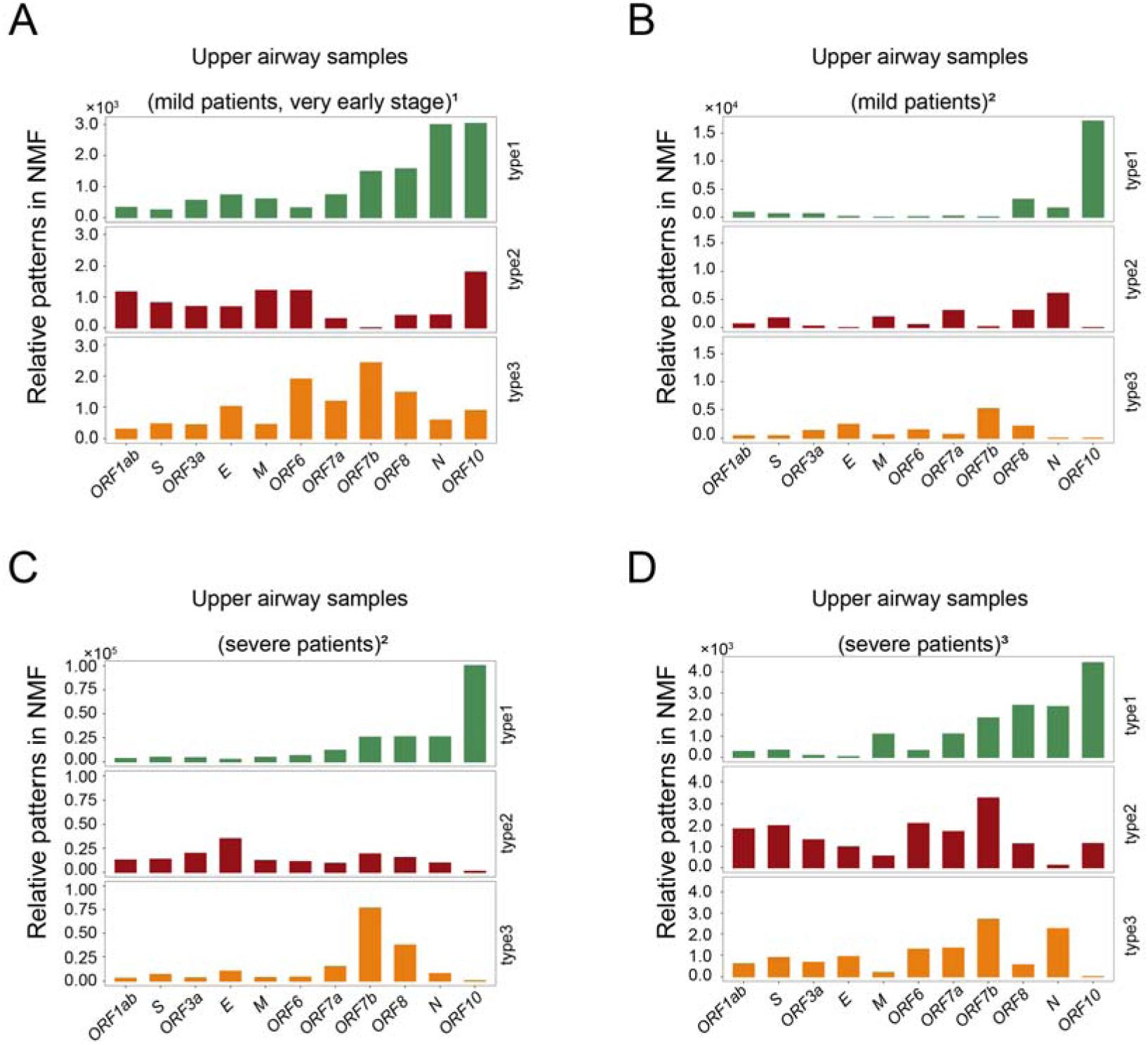
Unsupervised clustering identification of viral RNA expression patterns in metatranscriptome by using Non-negative Matrix Factorization (NMF). (A-D) Bar graph illustrating viral RNA transcription patterns in metatranscriptome samples computed by unsupervised machine learning. The type and source of the samples are labeled. 1: Data from Rajagopala, Seesandra V et al. 2: Data from Ren, Lili et al. 3: Data from Sulaiman, Imran et al. (The x-axis represents the genome of SARS-CoV-2, and the y-axis represents the relative abundance of each gene.).

Interestingly, this aligns well with biological expectations. In the results, we observed patterns with typical subgenomic transcription characteristics. Additionally, patterns with a relatively uniform distribution of viral RNA were noted, potentially representing viral particles released into the extracellular space after completing the viral life cycle within host cells. Notably, we identified a viral RNA expression pattern where only a portion of viral RNA had reads, and a significant amount of viral subgenomic RNA escaped detection by sequencing technology. This might indicate incomplete and deactivated viral fragments.

Therefore, we categorize the forms of SARS-CoV-2 shed into three types: viral replicons, viral particles, and viral fragments. "Viral replicon" refers to the intracellular form of SARS-CoV-2, where SARS-CoV-2 replicons and transcribes with a subgenomic manner. The transcription patterns of the SARS-CoV-2 genome vary among different target cells, so we categorize viral replicons based on cell types with viral tropism. Viral particles and viral fragments are defined as intact and fragmented SARS-CoV-2 particles, respectively.

### Supervised dissection of SARS-CoV-2 shedding patterns in metatranscriptomic data by using RedeCoronaVS

RedeCoronaVS is implemented as a linear programming model (Figure 4). It includes one objective function, several types of constraints, and several decision variables. The output of RedeCoronaVS is the abundance of the three viral shedding modes (Table S2). The number of decision variables is equal to the chosen input viral RNA detection rates for different cell types and expected viral RNA reads in the viral particle state, corresponding to the expectation matrix of constraints (STAR methods). The constraints of RedeCoronaVS ensure that the product of the expectation matrix input and the variables does not exceed the actual viral RNA expression. The objective function aims to maximize the abundance of viral replicons and viral particles to explain the observed SARS-CoV-2 gene expression data. The nonnegative constraints are used to confine the abundance of viral replicons and particles to be positive or at least zero, and the residuals between the observed gene expression and the estimated replicons and particles are used to infer the abundance of viral fragments. The subgenomic transcription pattern of SARS-CoV-2 is derived from published single-cell RNA sequencing data of SARS-CoV-2 infection (STAR Methods).

**Figure 4.**
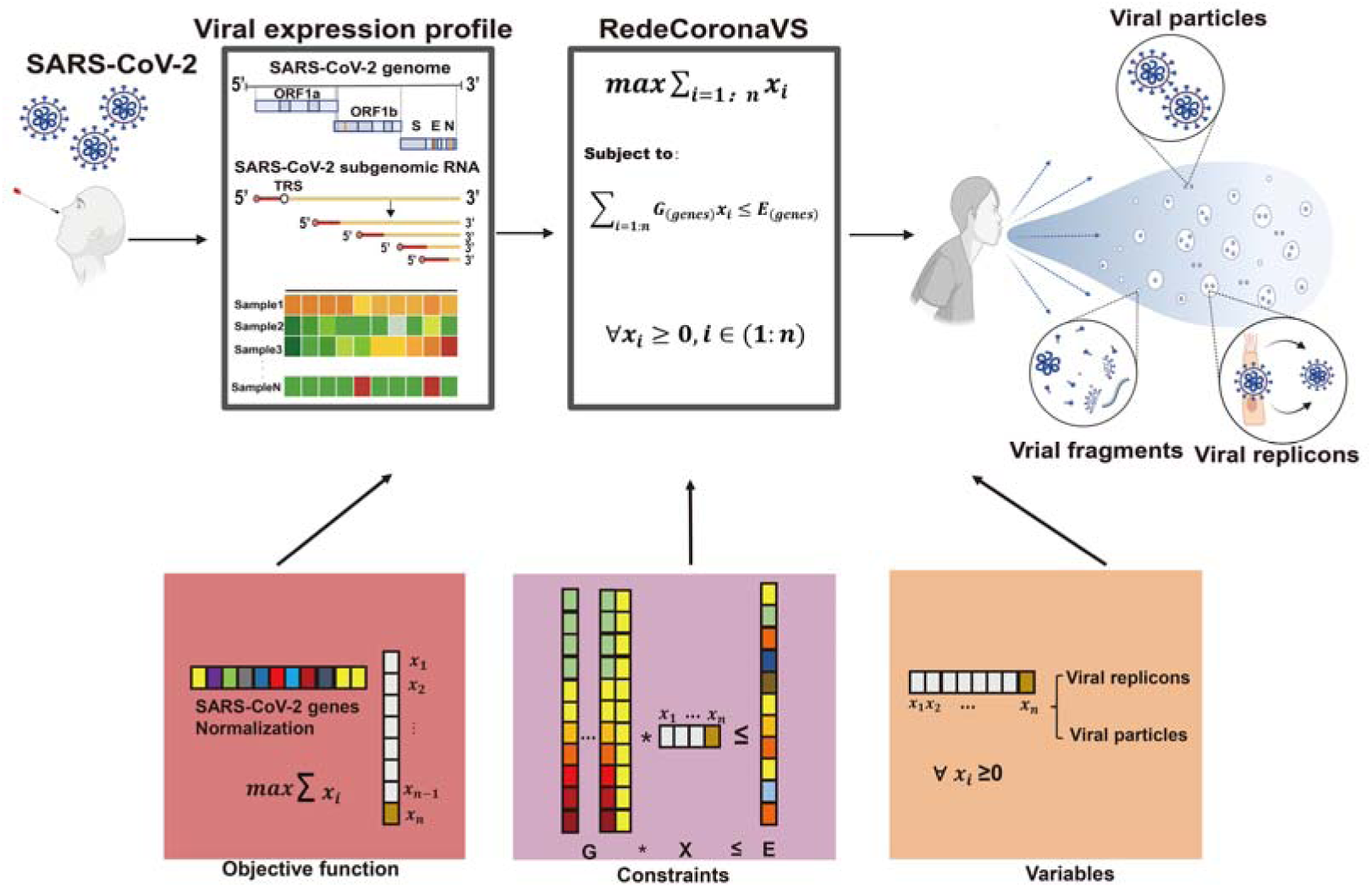
Workflow of RedeCoronaVS model. The viral RNA expression profile data, which was corrected for gene length, served as the input for the analysis. The RedeCoronaVS approach using linear programming was employed to determine the optimal fitting of viral replicons and viral particles in each sample. The residual values obtained from the model solution represented the presence of viral fragments. The final output provided information on the abundance of different viral shedding patterns. Created by BioRender.com.

To test the performance of RedeCoronaVS, we first conducted rigorous quality control on the samples to exclude the potential impact of sequencing depth (Figure S2). Next, to validate the accuracy of the RedeCoronaVS results, we conducted Pearson’s correlation analysis between the abundance of viral replicons/particles and CT values or viral RNA copies obtained from PCR test (Figure S3A-S3E). For ease of statistical analysis, we summed the total abundance of viral replicons. The analysis revealed a strong correlation between the abundance of viral replicons and viral particles with the PCR results in all cohorts except mild patients’ samples from the Wuhan cohort. This result is reasonable because mild cases in the later stages of the disease are expected to shed a large number of viral fragments. This could potentially yield positive nucleic acid test results without the presence of infectious viral replicons or particles being shed.

Subsequently, we aimed to validate the RedeCoronaVS results at a relative quantitative level for different viral shedding types. A fundamental assumption is that when COVID-19 patients primarily shed viral fragments as the main viral shedding type, it indicates that the patient has entered the late stages of infection. Simultaneously, corresponding PCR tests should detect little or no substantial amount of viral RNA^3^. To test this, we analyzed the relationship between the relative abundance of the three viral shedding types in different samples and PCR results (STAR Methods). The ternary plot results confirm our hypothesis: when the proportion of viral fragments is higher, there is a tendency to detect a lower amount of viral RNA (Figure S4A-S4C). These results indicate that RedeCoronaVS accurately differentiates and precisely quantifies viral shedding patterns in respiratory tract of COVID-19 patients.

### RedeCoronaVS revealed relationships between different viral shedding patterns

We then explored correlations between viral replicons, viral particles and viral fragments. We calculated the relative quantification of each mode for each sample and conducted correlation analysis (Figure 5A-5C). The results showed a positive correlation between viral replicons and viral particles, consistent with previous studies.^21^ Viral fragments exhibited a strong negative correlation with the other two modes which indicates that the proportion of viral fragments can serve as a recovery indicator for COVID-19 patients. Interestingly, in the Vanderbilt cohort, providing samples from the early stages of infection, we observed negative correlations among all three viral shedding modes (Figure 5A).

**Figure 5.**
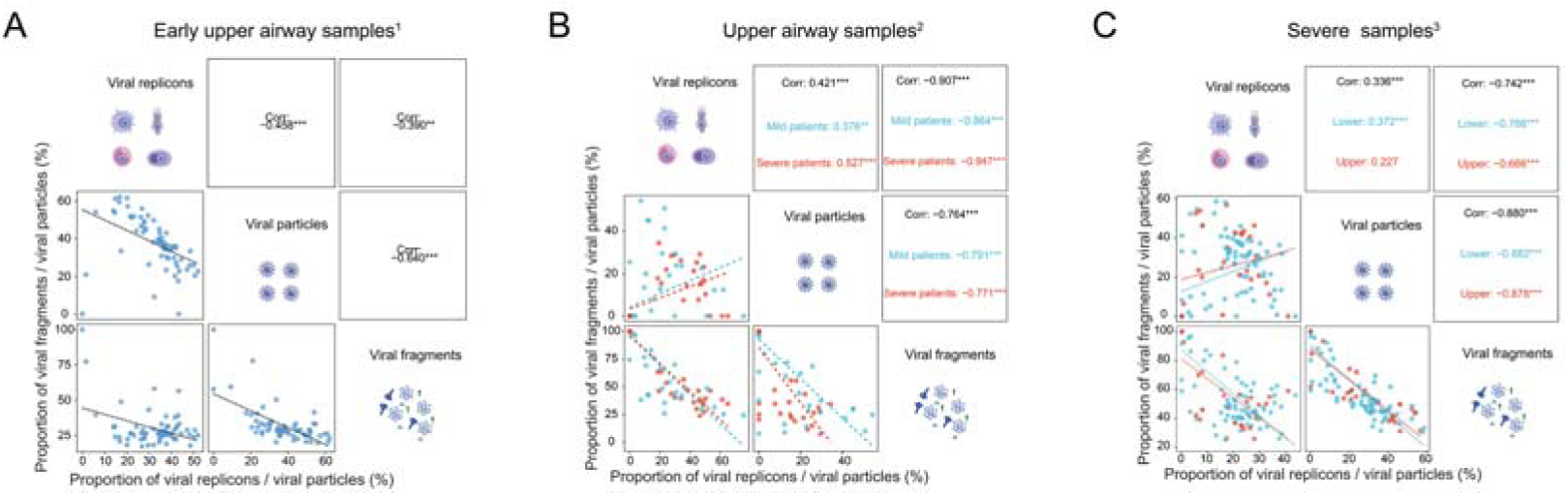
Correlations between different viral shedding patterns. Correlations between viral replicons, viral particles and viral fragments. (A-C) Plot of the relationship between the relative abundance of different types of viral shedding patterns. (Pearson correlation, dotted line) Asterisks indicate significant correlations at P < 0.05 (*), P < 0.01 (**) or P < 0.001 (***). The type and source of the samples are labeled. 1: Data from Rajagopala, Seesandra V et al. 2: Data from Ren, Lili et al. 3: Data from Sulaiman, Imran et al.

### Viral replicons are associated with clinical status of COVID-19 patients

Next, a cluster analysis was conducted on the RedeCoronaVS results obtained. Consistent with the fact that most samples were collected within the recovery phase, the majority of samples exhibited high levels of viral fragments but low levels of viral replicons or particles (Figure 6A). The large cohort size enabled us to dissect the associations of sampling site, days after hospitalization and COVID-19 severity with the abundance of different viral shedding patterns in patients. We applied ANOVA analysis to interrogate such associations based on 203 upper airway samples and 95 lower airway samples (Figure 6B; Table S3; STAR Methods). Indeed, we have observed variations in viral replicons types in respiratory shed across different clinical stages of COVID-19 patients. The proportion of replicons in neutrophils and macrophages is closely correlated with the sampling time after hospitalization (Figure 6C-6D). It is significantly higher when patients are admitted early compared to one week after admission, indicating that SARS-CoV-2 primarily sheds innate immune cells during the early stages of infection. The proportion of replicons in macrophages is associated with the sampling site, being more commonly detected in upper respiratory tract samples (Figure 6E). Replicons in plasma cells are more likely to be detected in severe COVID-19 patients, suggesting that they may be the primary cell type shedding in this category of patients (Figure 6F). In summary, viral replicons are closely related with the clinical status of the COVID-19 patients.

**Figure 6.**
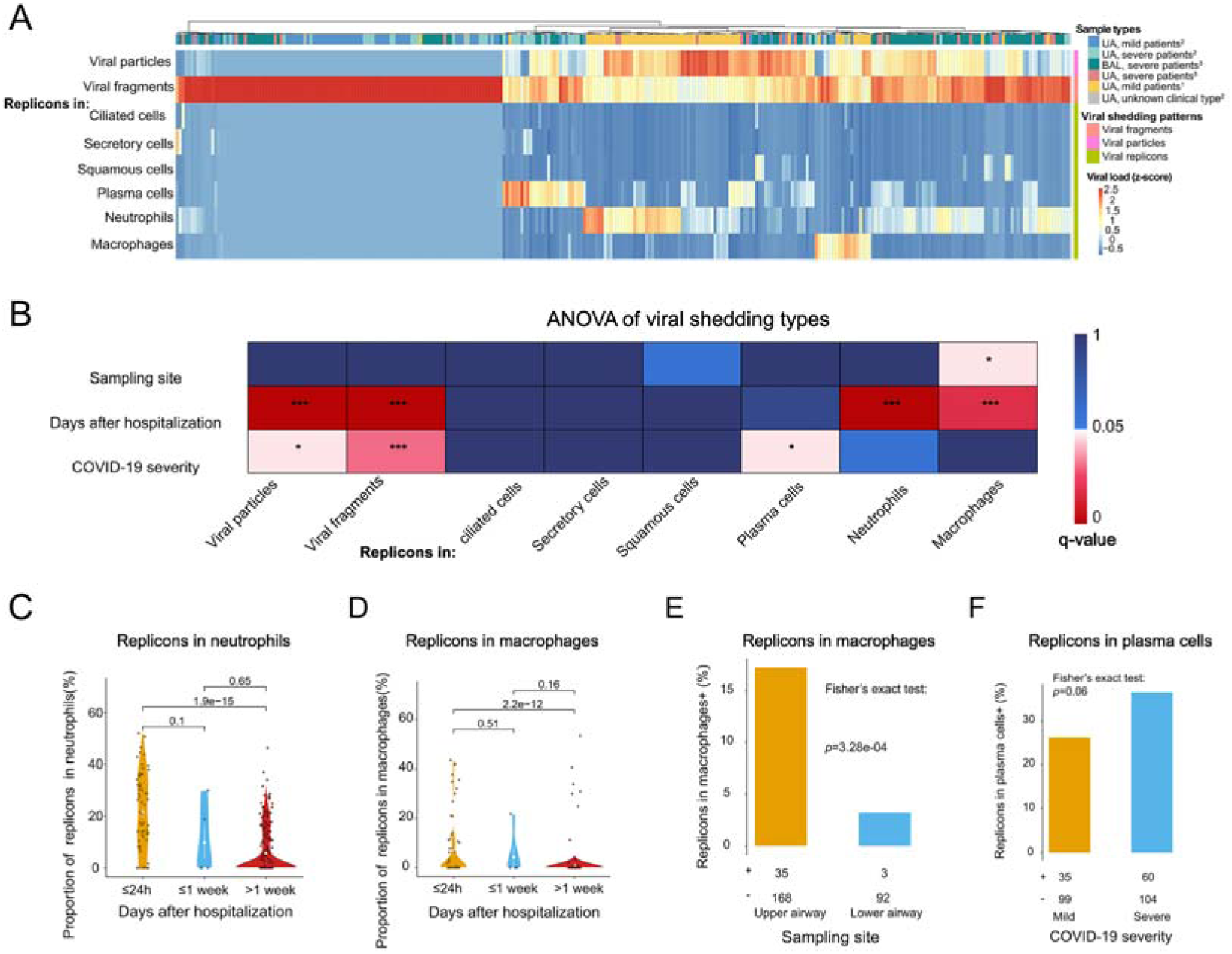
Correlations between viral shedding types and clinical data. (A) Heatmap of overall viral shedding in COVID-19 patients. UA: Upper airway samples. BAL: Lower airway samples. (B) Heatmap for q values of ANOVA. Asterisks indicate significant correlations at q < 0.05 (*), q < 0.01 (**) or q < 0.001 (***). (C) Violin plot of the relative abundance of replicons in neutrophils changing with the hospitalization time of COVID-19 patients. (D) Violin plot of the relative abundance of replicons in macrophages changing with the hospitalization time of COVID-19 patients. (E) Replicons in macrophages are more likely to be detected in upper respiratory tract samples. (F) Replicons in plasma cells are more likely to be detected in severe COVID-19 patients.

### Associations of SARS-CoV-2 shedding patterns with sampling time and COVID-19 severity

Lastly, we aimed to identify the shedding phases of SARS-CoV-2 and other potential factors influencing viral shedding by RedeCoronaVS. ANOVA revealed that viral replicons, viral particles, and viral fragments are all heterogeneously associated with factors such as sampling days after hospitalization and disease severity (Figure 6B). Therefore, we focus on analyzing individual cohorts to disregard the influence of other potential errors. As the Wuhan cohort provided the clinical information of COVID-19 patients, i.e., days from patient hospitalization to sample collection and patients’ clinical status, we applied Fisher’s exact test to evaluate whether significant changes exist between the first week of hospitalization and later (Figure 7A). The result indicates that COVID-19 patients are highly infectious within the first week of hospitalization and the infectiousness decreased later. Consistently, upon further analysis of the changes in the proportion of viral replicons and viral fragments in total viral RNA, we observed a significant change in the proportion of viral replicons and viral fragments shed by COVID-19 patients one week after hospitalization (Figure 7B-7C). Patients tested early in hospitalization exhibit a higher proportion of viral replicons and a lower proportion of viral fragments compared to those tested later. In another independent cohort, we observed that early hospitalization COVID-19 patients primarily shed viral replicons and particles, with their proportions significantly higher than viral fragments, consistent with the results from the Wuhan cohort (Figure 7D). Notably, we observed that patients predominantly shed viral particles in the early stages of viral infection (Figure 7D). Our findings provide molecular supports to the notion that COVID-19 patients are highly contagious during the initial week of hospitalization, and highlight the importance of early isolation measures for individuals displaying symptoms of COVID-19.^22^

**Figure 7.**
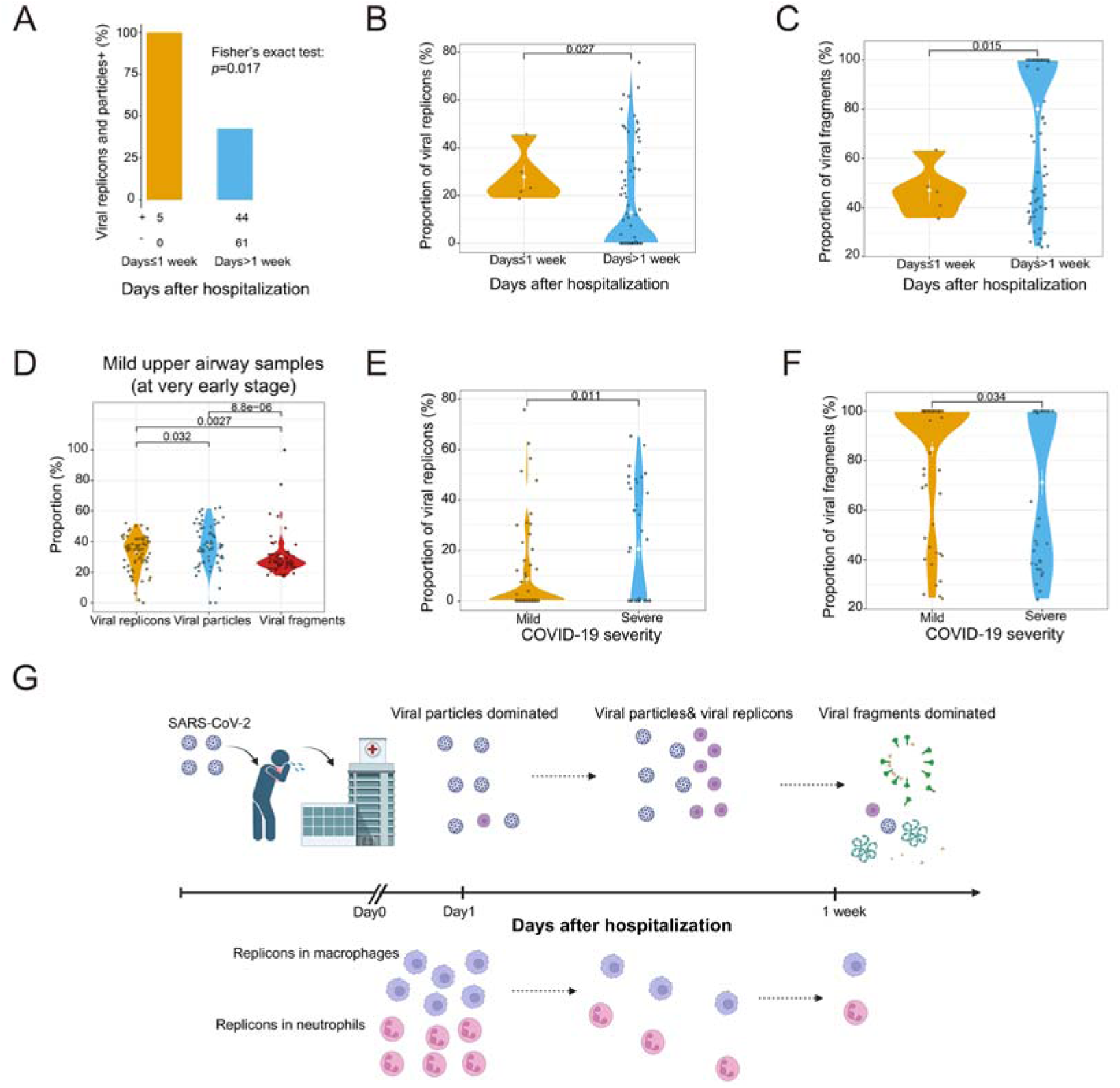
Biological discoveries from RedeCoronaVS. (A) Fisher’s exact test revealed a significant clustering of viral shedding within 1 week of SARS-CoV-2 infection. (B) Violin plot of the difference in the proportion of viral replicons in the total viral RNA read count within one week and after one week of symptom onset in COVID-19 patients. (Wilcox’s rank test, the mean and standard error of the samples are indicated.). (C) Violin plot of the difference in the proportion of viral fragments in the total viral RNA read count within one week and after one week of symptom onset in COVID-19 patients. (Wilcox’s rank test, the mean and standard error of the samples are indicated.) (D) Violin plot comparing the relative abundance of shedding patterns in early-stage COVID-19 patients. (Data from Rajagopala, Seesandra V et al.). (E) Violin plot illustrating the difference in the proportion of viral replicons in the total viral RNA read count between mild and severe COVID-19 patients. (F) Violin plot of the difference in the proportion of viral fragments in the total viral RNA read count between mild and severe COVID-19 patients. (G) The diagram illustrates the patterns of viral shedding in COVID-19 patients at various stages of infection. Created by BioRender.com.

Whether there is a difference in viral shedding between mild and severe COVID-19 patients is controversial.^23,24^ To address this question, we compared the relative abundance of viral replicons and fragments in mild and severe patients using the Wuhan cohort (Figure 7E-7F). We excluded samples within the first week of hospitalization to avoid potential time-related effects. We observed a significant variation in the relative abundance of viral replicons and fragments in severe patients compared to mild patients. Compared to mild patients, the high proportion of viral replicons and low proportion of viral fragments in severe COVID-19 patients suggest that they may continue to shed infectious viral replicons in the late stages of infection. The underlying cause for this difference may be the immunological dysregulation in severe patients, preventing the complete clearance of the virus from the body.^25^ Therefore, our results indicate that the severity of the disease is also a crucial factor influencing viral shedding.

Based on these results, we summarized the pattern and type of viral shedding in COVID-19 patients at different time points (Figure 7G). Viral particle shedding dominated in the early stages of hospitalization, transitioning to viral replicons and particles as the disease progressed. One week after hospitalization, patients predominantly shed viral fragments as their infectivity is significantly reduced.

## Discussion

Understanding the biological characteristics of SARS-CoV-2 shedding in the respiratory tract is crucial for formulating more precise control policies in the future.^3^ Despite extensive research on the abundance and characteristics of SARS-CoV-2 shedding, determining the infectivity of specific COVID-19 patients remains challenging. While viral load is commonly used as a surrogate for infectious viral particles, it only indicates the presence of viral RNA in the sample, not infectivity. The isolation and quantification of live virus require specialized personnel, equipment, and facilities, as well as strict sample preservation and transportation protocols, making it impractical for studying viral shedding in large patient cohorts.^3^ Therefore, there is an urgent need for new methods to provide innovative directions for studying viral shedding issues.

High-throughput sequencing of viral RNA provides a potential solution, offering quantitative information on the entire viral transcriptome with advantages such as simplicity and cost-effectiveness. Yet, whole transcriptome sequencing data is complex and requires specific tools for precise decoding of shedding patterns. This study establishes the correlation between metatranscriptomic data from respiratory samples of COVID-19 patients and SARS-CoV-2 viral shedding. Unsupervised machine learning methods help identify potential viral shedding patterns. Building upon this, we introduce a tool called RedeCoronaVS. Developed based on linear programming models, RedeCoronaVS achieves deconvolution of sequencing data by incorporating pre-existing information to estimate the abundance of different viral shedding patterns. Furthermore, RedeCoronaVS enhances our understanding of viral shedding in COVID-19 patients. Viral replicons are correlated with the sampling time and clinical status of COVID-19 patients. Specific types of viral replicons may represent critical clinical information, and they tend to co-shed with viral particles. Our results indicate that patients shed a significant amount of infectious virus within the first week of hospitalization. In severe cases, patients may continue shedding the virus during the late stages of infection, suggesting disease severity is an important factor affecting viral shedding.

In summary, our study presents RedeCoronaVS as a valuable tool for detecting and quantifying viral shedding patterns in COVID-19 patients. By analyzing metatranscriptomic sequencing data of SARS-CoV-2 from hundreds of COVID-19 respiratory samples, RedeCoronaVS enables the identification of the specific types and abundance of viral presence, thereby providing insights into the disease progression in patients. Importantly, RedeCoronaVS is highly scalable and holds potential for application in other coronavirus diseases, making it a valuable tool for future responses to emerging respiratory infectious diseases.

### Limitations of the study

Due to the challenges in directly dissecting intact viral particles, viral replicons within host cells, and viral fragments from respiratory specimens of large cohorts of COVID-19 patients, it is difficult to validate our results with current experimental protocols. Another limitation is the absence of information regarding the precise onset time of COVID-19 in patients. This study relies on the patients’ hospitalization records to estimate the real onset time, potentially leading to underestimation and uncertainty.

## STAR METHODS

### KEY RESOURCES TABLE

| REAGENT or<br>RESOURCE | SOURCE | IDENTIFIER |
| --- | --- | --- |
| **Software and Algorithms** |  |  |
| R | N/A | https://www.cran.r-project.org |
| R Studio | N/A | https://www.rstudio.com |
| STAR | Dobin,<br>Alexander et<br>al., 2012<sup>26</sup> | https://github.com/alexdobin/STAR |
| featureCounts | Liao, Yang et<br>al., 2013<sup>27</sup> | https://sourceforge.net/projects/subread/ |
| ggtern (R package) | Nicholas E.<br>Hamilton et<br>al., 2018<sup>28</sup> | http://www.ggtern.com/d/2.2.2/ |
| NMF (R package) | Brunet JP, et<br>al., 2004<sup>29</sup> | https://cran.r-project.org/web/packages/NMF/index.html |
| GGally | N/A | https://cran.r-project.org/web/packages/GGally/index.html |
| RedeCoronaVS | This paper | https://github.com/spencerJin/RedeCoronaVS |
| **Deposited data** |  |  |
| Sing-cell<br>SARS-CoV-2 viral<br>RNA detection rate | Ren, Xianwen<br>et al., 2021<sup>16</sup> | N/A |
| metatranscriptomic<br>data for viral RNA<br>reads | Ren, Lili et<br>al., 2021<sup>30</sup> | N/A |
|  | Sulaiman,<br>Imran et al.<sup>31</sup> | SRA project number: PRJNA688510 |
|  | ajagopala,<br>Seesandra V<br>et al. | SRA project number: PRJNA922078 |

### RESOURCE AVAILABILITY

#### Lead Contact

Further information and requests for resources and reagents should be directed to and will be fulfilled by the Lead Contact, Jianwei Wang.

#### Materials Availability

This study did not generate new unique reagents.

#### Data and Code Availability

The samples from the Wuhan cohort were published in the publication by Ren, et al., doi: 10.1164/rccm.202103-0814OC.30 As the original data was not publicly available, we requested the raw data from the corresponding author. The NYU Manhattan Bronchoscopy Cohort data are deposited in the Sequence Read Archive (SRA) database under the National Center for Biotechnology Information (NCBI), with project number PRJNA688510. The Vanderbilt University Cohort data are also available in the SRA database under project number PRJNA922078. The single-cell level SARS-CoV-2 viral detection rate data were obtained from the supplementary information of Ren et al.’s publication.^16^ RedeCoronaVS is available at https://github.com/spencerJin/RedeCoronaVS.

## Method Details

### Data collection and processing

The metatranscriptomic data used in this study were publicly available. The Wuhan Cohort data were published in Ren, et al, which contained 634 oropharyngeal swab metatranscriptome sequencing data from COVID-19 patients.^30^ In the Wuhan cohort, we defined mild or severe COVID-19 patients based on previous research.^32^ The criteria for this definition were whether patients required interventions such as supplemental oxygen, mechanical ventilation, or ECMO. Patients requiring such treatments were classified as severe COVID-19 patients, while those not requiring treatment were classified as mild COVID-19 patients. The NYU Manhattan Bronchoscopy Cohort and the Vanderbilt University Cohort were downloaded from NCBI’s SRA database under project numbers PRJNA688510 and PRJNA922078, respectively.^31,33^ We gathered 182 samples including 64 upper airway samples and 118 lower airway samples in the NYU Manhattan Bronchoscopy Cohort and 67 upper airway samples in the Vanderbilt University Cohort.

The samples’ raw data are kept in FASTQ format, and we employ the same technique for upstream processing. All data were mapped to the SARS-CoV-2 genome NC_045512 using STAR with default parameters.^26^ Gene counts were summarized using the featureCounts program.^27^ We normalized the viral gene counts for each sample based on viral gene length (viral gene counts/viral gene length) to create the viral gene expression profile for each sample to prevent the effect of gene length on abundance for viral gene reads. Furthermore, we applied a filtering process to the samples based on sequencing depth to mitigate the impact of low sequencing depth on the accuracy of our results. Finally, A total of 301 samples from the three cohorts were included in the analysis.

### Using Non-negative Matrix Factorization (NMF) method identifies potential viral shedding patterns

The general formulation of non-negative matrix factorization (NMF) is as follows: Given a non-negative matrix X∈R^n*m^ and a user-specified K≤n, the objective is to provide a low-rank approximation matrix. To identify the viral shedding patterns of SARS-CoV-2, we utilize NMF to decompose the count matrix (E) of viral genes for N genes and M samples into two matrices, W and H, such that E∼WH^T^. The amplitude matrix W has columns associated with features of the viral genes, and the pattern matrix H has rows associated with features of the samples. The matrix W represents different types of viral shedding modes. The calculations for NMF in this study were conducted using the R package NMF.^29^

### Modeling SARS-CoV-2 viral shedding patterns in metatranscriptome samples

We developed RedeCoronaVS based on linear programming to estimate the viral shedding patterns from the viral RNA expression of the samples. Here, we propose a hypothesis regarding the shedding of the virus in the respiratory tract of COVID-19 patients. We suggest that there are three modes in which the virus can be detected: active viral replication within target cells (viral replicons), free viruses released after completing their life cycle (viral particles), and inactive viral fragments (viral fragments). Each of these modes is associated with distinct patterns of viral RNA expression. In addition, after normalizing for gene length, the expression of individual viral RNAs within viral particles is expected to be homogeneous. Based on the aforementioned assumptions, we constructed an expectation matrix consisting of two parts. The first part represents the viral RNA expression information within SARS-CoV-2 infected cells obtained through single-cell sequencing from previous research.^16^ The second part represents the simulated average expression of viral RNA, which serves to characterize the viral particles. Considering that incomplete viral RNA expression is thought to be present in the sample, viral fragments can be represented by the residuals produced by RedeCoronaVS.

### 1.1 Notations

G denotes the expectation matrix, which was defined previously as the expected expression pattern of SARS-CoV-2 RNA under specific conditions. E denotes normalized real viral RNA sequencing data. We use (x_1_, x_2…_ x_n_) as decision variables, and these variables represent the abundance of viral replicons and viral particles, respectively. The number of decision variables depends on the number of columns in the G matrix.

### 1.2 Linear programming model

We formulate the pattern of viral shedding problem as an optimization problem using the linear programming model. The objective is to maximize the sum of decision variables to obtain the best fit to the observation matrix.

Since we have normalized the observation matrix such that the sum of each column is 1, the objective function is:

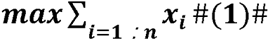

Let genes= (*ORF1ab, S, ORF3a, E, M, ORF6, ORF7a, ORF7b, ORF8, N, ORF10*), therefore, the constraints should be satisfied that the fitted value of each gene does not exceed the actual observed value of the sample. The constraints are subject to:

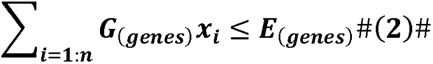

The decision variables should satisfy the following conditions:

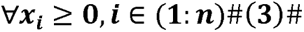

### Statistical analysis

To evaluate the impact of different clinical statuses of COVID-19 patients on the characteristics of viral shedding, we conducted multivariate ANOVA analysis on the proportions of viral replicons within different cells, viral particles, and viral fragments, along with clinical information including sampling sites, sampling time after hospitalization, and disease severity. Multiple testing corrections were performed, and results with FDR < 0.05 were considered significant.

Fisher’s exact test was used for discovering significant viral shedding time points. Pearson correlation was used for correlation analysis. Results with statistical significance should have a p-value less than 0.05. The ternary plots were created using package ggtern.^28^ All statistical analysis was implemented in R (4.2.1).

## ACKNOWLEDGMENTS

This study was funded by Science Fund for Creative Research Groups of the National Natural Science Foundation of China (82221004), the National Natural Science Foundation (81930063), the Chinese Academy of Medical Sciences Innovation Fund for Medical Sciences (2021-I2M-1–038) and Changping Laboratory research fund.

## AUTHOR CONTRIBUTIONS

X.R. conceived the project. L.R., X.R., and J.W. supervised the project. X.R. and X.J. developed RedeCoronaVS algorithm. X.J. performed all of the analysis and wrote the paper. L.R., X.R., and J.W. edited the manuscript.

## Supplemental information

### Integrative single-cell and metagenomic analysis dissects SARS-CoV-2 shedding modes in human respiratory tract

**Figure S1.**
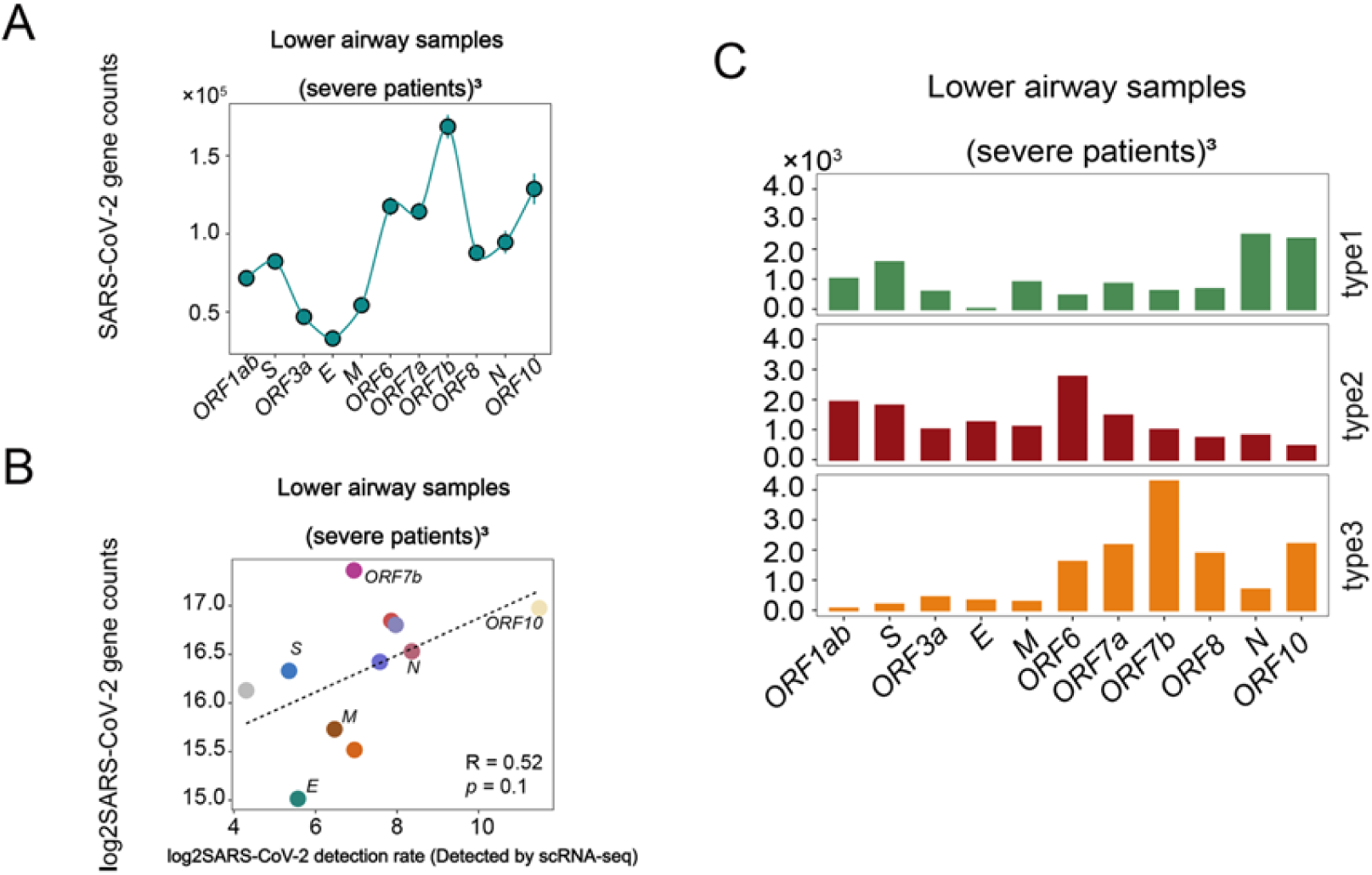
Results of lower airway samples. (A) Line graphs depicting the mean distribution of various viral genes in metatranscriptome samples. (Viral gene reads were normalized by TPM; standard error of the samples is indicated.). (B) Plots of the relationship between the normalized mean expression of SARS-CoV-2 viral genes in lower airway metatranscriptome samples and the detection rate of SARS-CoV-2 in ciliated cells. (Pearson correlation; dotted line, the logarithmic transformation was applied to viral detection rate.). (C) Bar graph illustrating viral RNA transcription patterns in metatranscriptome samples computed by unsupervised machine learning. Data from Sulaiman, Imran et al. (The x-axis represents the genome of SARS-CoV-2, and the y-axis represents the relative abundance of each gene.). Data from Sulaiman, Imran et al.

**Figure S2.**
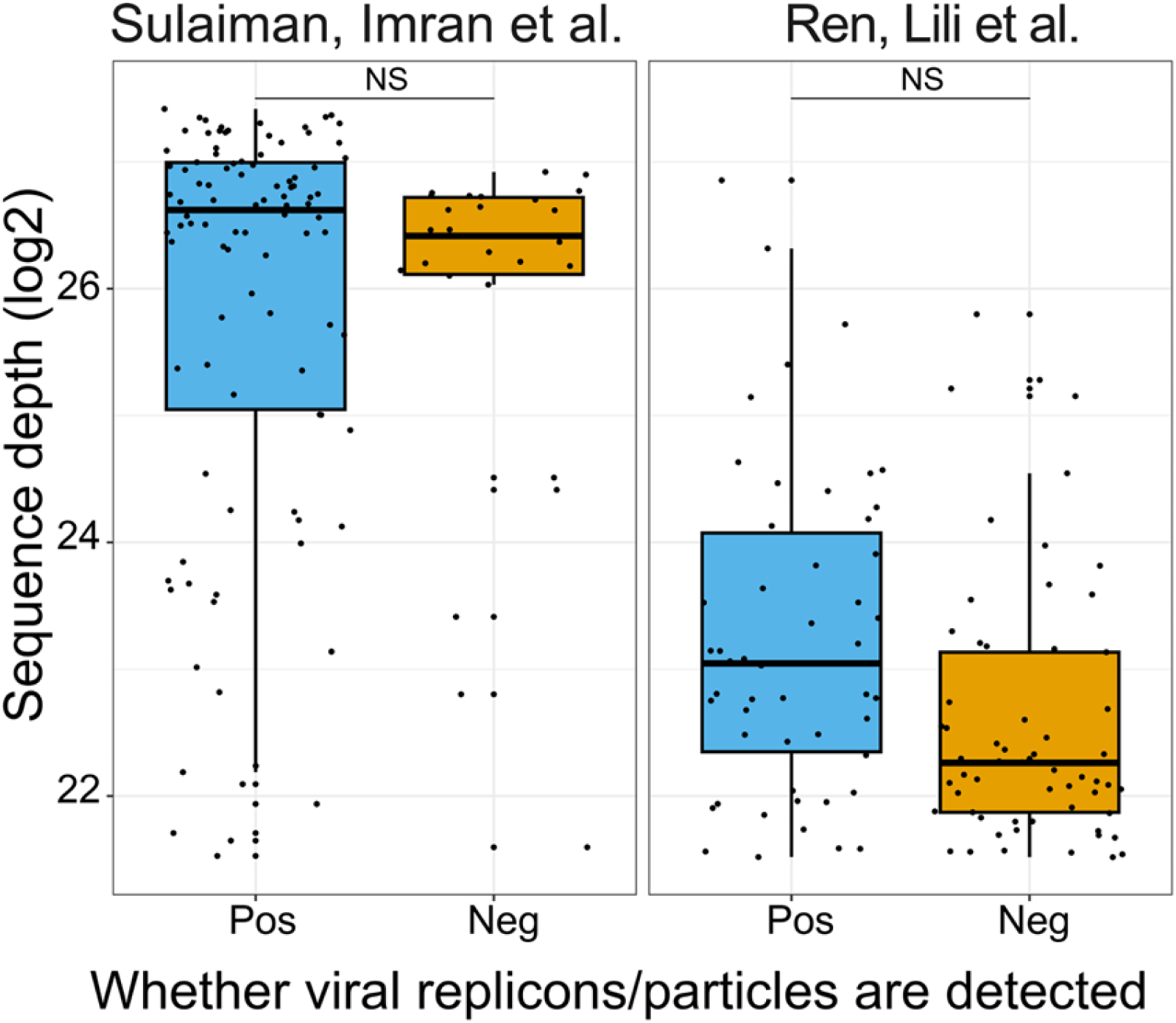
Excluding the impact of sequencing depth on the VSD model. A strict sequencing depth threshold was adopted to eliminate potential sequencing depth differences among different viral detection patterns. (Threshold=3000000)

**Figure S3.**
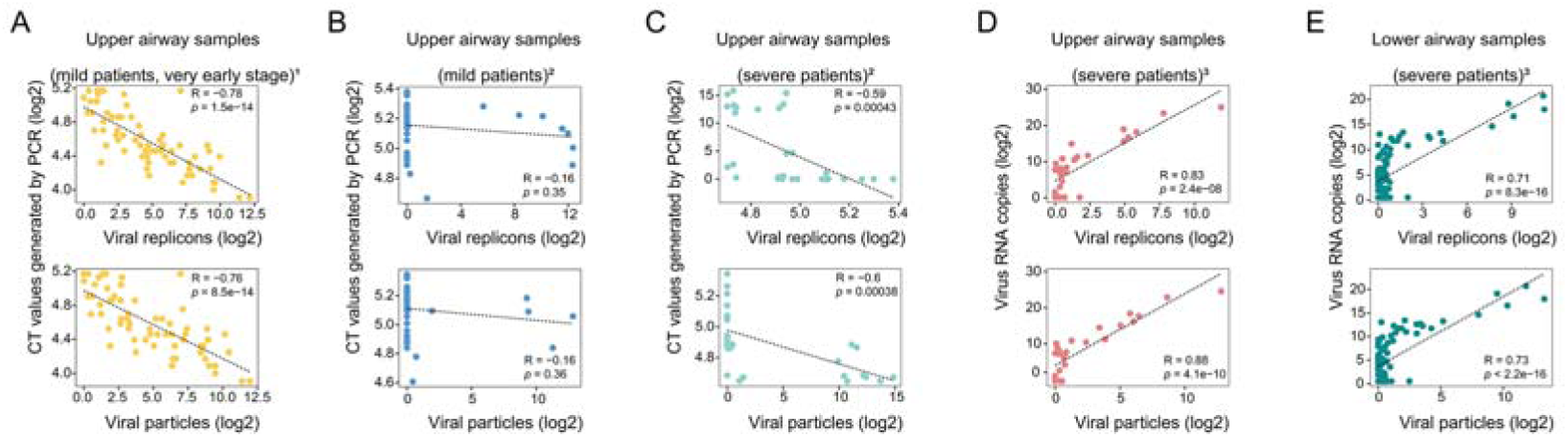
Validations of RedeCoronaVS. (A-E) Plot of the relationship between the abundance of two viral shedding patterns and the results obtained from RT-qPCR (CT values or viral RNA copies) which demonstrates a strong correlation between the shedding abundance and the viral load. (Pearson correlation, dotted line.). The type and source of the samples are labeled. 1: Data from Rajagopala, Seesandra V et al. 2: Data from Ren, Lili et al. 3: Data from Sulaiman, Imran et al.

**Figure S4.**
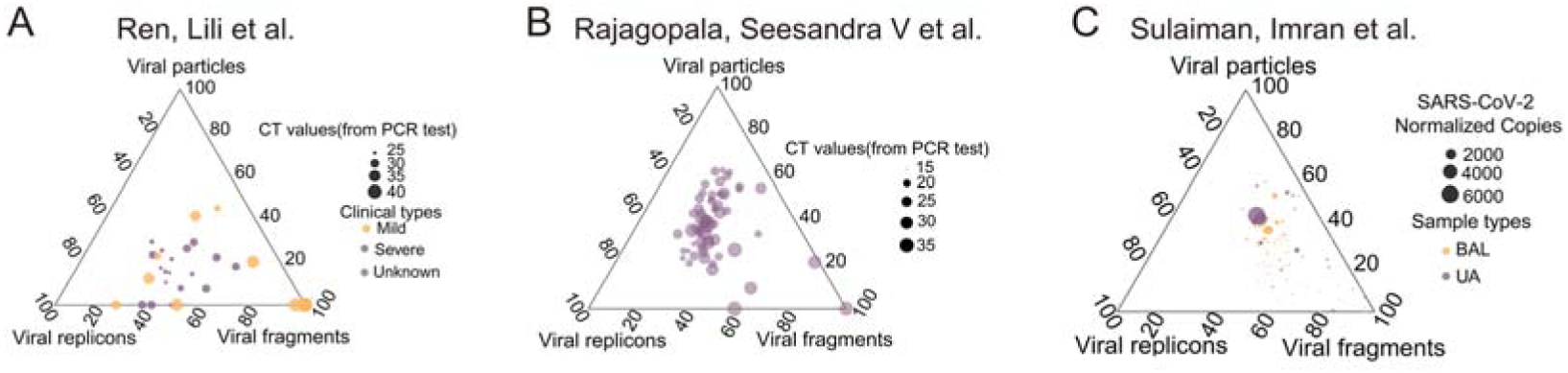
Ternary plots showing the relationship between relative abundance and viral load for different viral shedding patterns. (A-C) Ternary plots across 3 types of viral shedding patterns. Each point’s size corresponds to the CT value or viral RNA copy number of the sample, and its position is determined by the contribution of the relative abundance of different shedding patterns in the sample. The type and source of the samples are labeled.

