## Supplementary figures and images for "SARS-CoV-2 shedding dynamics in human respiratory tract"

### Supplemental Figure 1

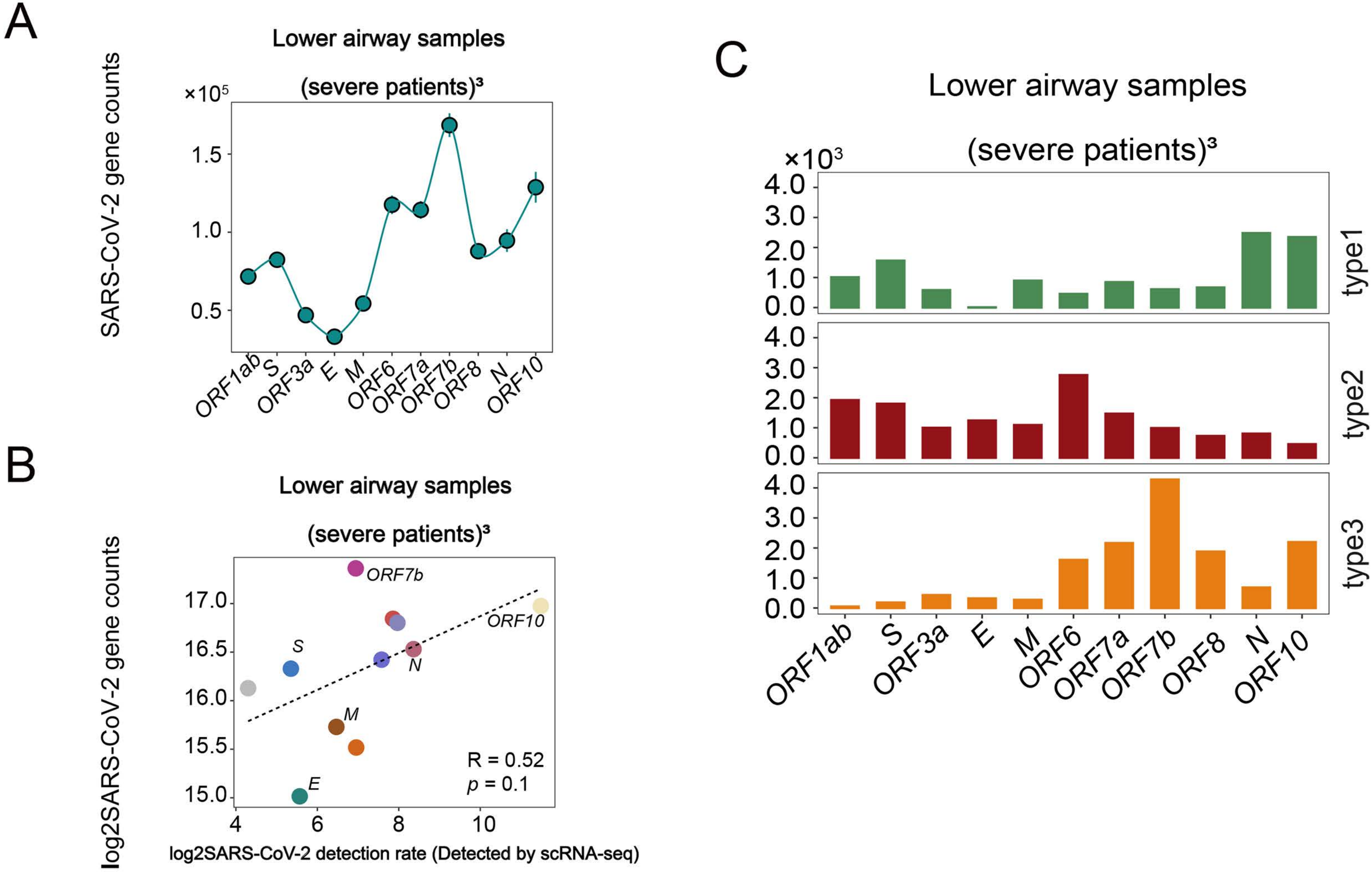

### Supplemental Figure 2

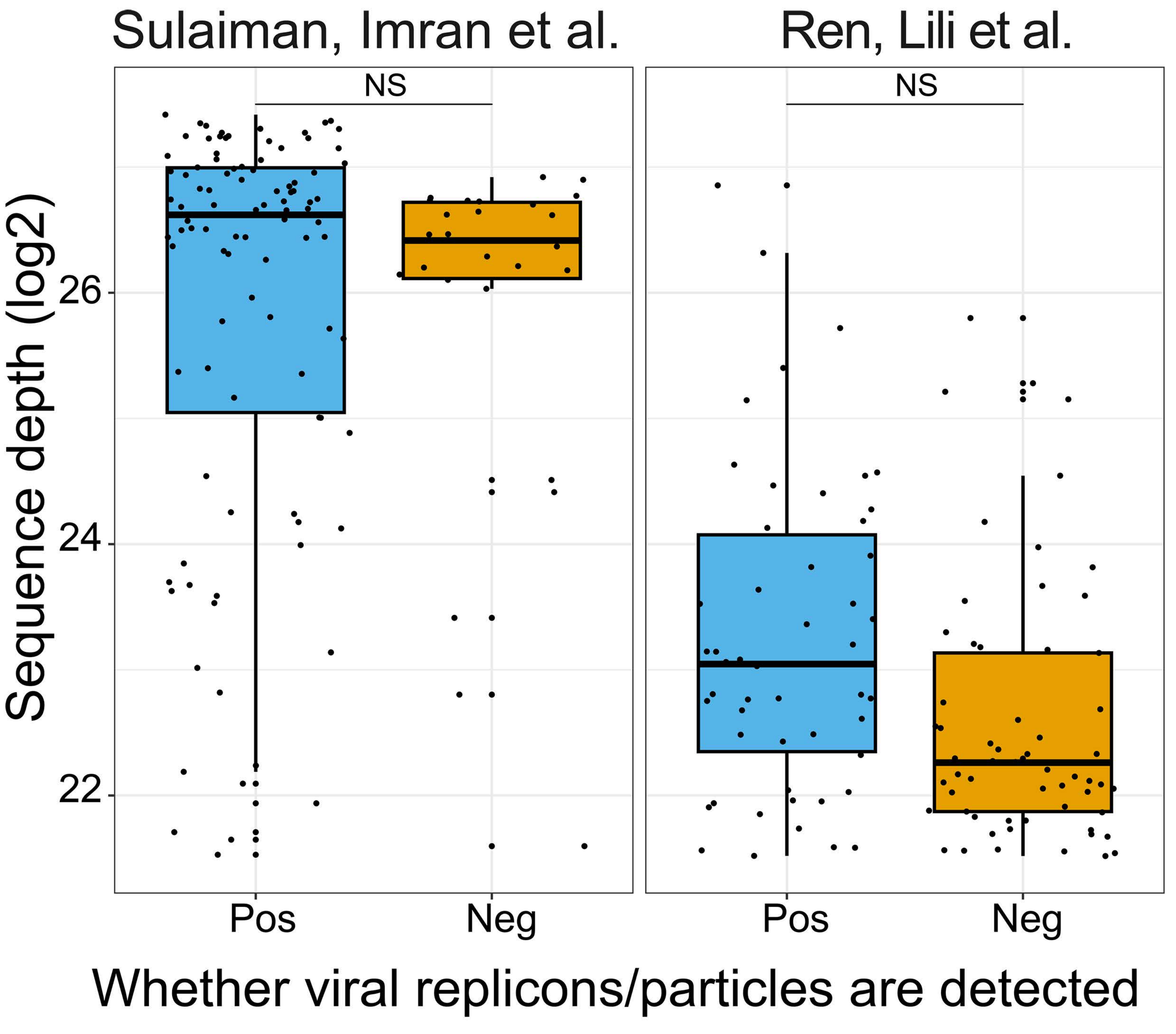

### Supplemental Figure 3

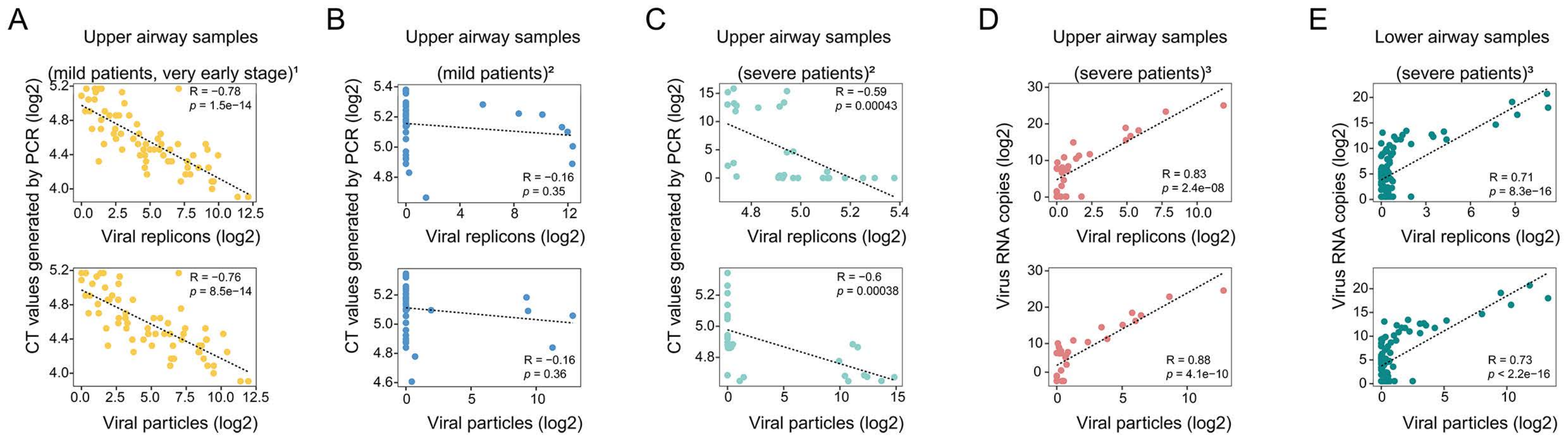

### Supplemental Figure 4

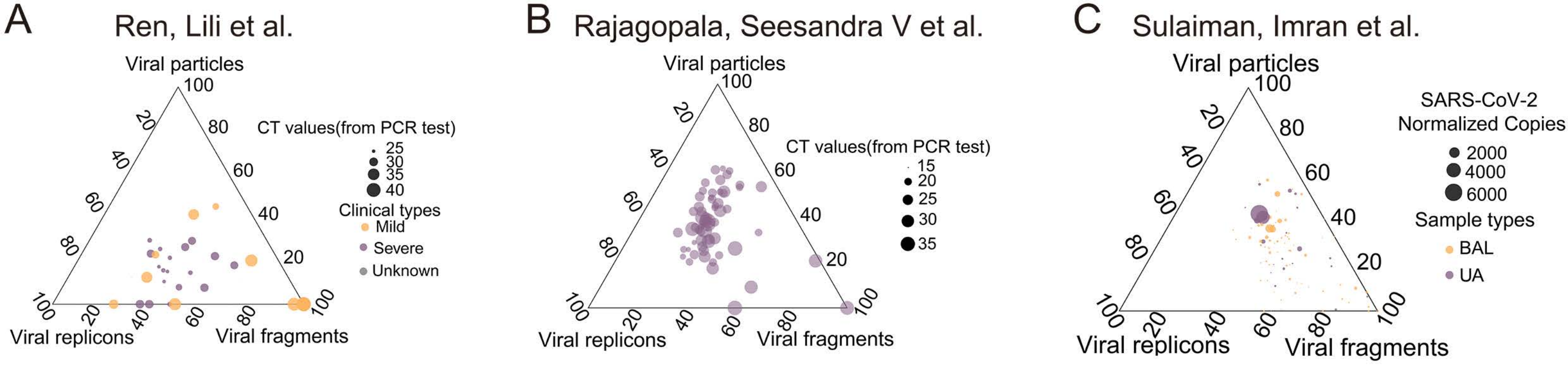
